# The heat sensitivity of sperm in the lizard *Anolis sagrei*

**DOI:** 10.1101/2024.08.14.607821

**Authors:** Wayne Wen-Yeu Wang, Natalie R. Page, Anthony M. Strickler, Alicia K. Kusaka, Alex R. Gunderson

## Abstract

The heat sensitivity of reproduction is a critical determinant of population persistence under climate change. However, the heat sensitivity of gametes is poorly known relative to adults. We developed a method to measure the heat tolerance of lizard sperm cells, and used the method to test several aspects of sperm cell thermal biology in the brown anole lizard (*Anolis sagrei*). We estimated the repeatability of sperm traits by measuring heat tolerance and baseline motility of ejaculated sperm from the same individuals multiple times over 21 days. To investigate co-adaptation of sperm and adult thermal traits, we tested for a correlation between sperm heat tolerance and the heat tolerance of adults that produced them. Furthermore, we tested for effects of episodic heat stress experienced by males on sperm performance. Sperm heat tolerance and motility were both repeatable, consistent with evolutionary potential, though there was clear evidence for environmental effects on these traits as well. Contrary to the expectation of thermal co-adaptation, we found no correlation between sperm and adult heat tolerance. A single, episodic extreme heat event experienced by adult males immediately impaired sperm motility, consistent with detrimental effects of adult heat stress on stored sperm. Our study adds to the mounting evidence that sperm are heat-sensitive and represent a vulnerability to global warming, but also suggest evolutionary potential for thermal adaptation at the gamete level.

**Summary statement:** This study investigates gamete heat sensitivity in lizards, revealing heat tolerance and repeatability in sperm thermal traits. These findings are essential for understanding reproductive responses to climate change.

## Introduction

Global climate change is making thermal environments increasingly unpredictable and extreme (IPCC, 2014). As a result, understanding how organism will respond to challenging new temperatures and predicting which taxa will be most vulnerable to change is a major goal for biologists (Williams et al., 2008; Huey et al., 2012; Hughes, 2000). Most research on warming in animals is done through the lens of adult or juvenile thermal tolerance based on the measurement of traits like critical thermal maximum (CTmax) or lethal temperature (Pinsky et al., 2019; Buckley and Kingsolver, 2021; Gunderson and Stillman, 2015). The thermal tolerance of haploid gametes such as sperm is much less studied despite their critical role in reproduction and population persistence (Dougherty et al., 2024; Reinhardt et al., 2015; Walsh et al., 2019; Wang and Gunderson 2022).

There is increasing evidence that heat decreases sperm quality and fertilization success (Wang and Gunderson, 2022; Walsh et al., 2019; Dougherty et al., 2024). For example, male flour beetles (*Tribolium castaneum*) produce significantly fewer viable sperm and sire fewer offspring after experiencing heat wave conditions (Sales et al., 2018, 2021; Vasudeva et al., 2021). Similarly, sperm performance and fertilization success in multiple sea urchin species decreases when ejaculated sperm are exposed to heat (Rahman et al., 2009; Binet and Doyle, 2013). In a larger ecological context, the sterility temperatures of male *Drosophila* better predict species distributions and extinction risk than either female sterility temperature or adult heat limits (Parratt et al., 2021; van Heerwaarden and Sgrò, 2021). These finding demonstrate the importance of male gametes in dictating responses to global warming and highlight the need to better understand sperm thermal biology. This is especially true in reptiles, for which there is a lack of data on the heat sensitivity of reproduction relative to other systems (Dougherty et al., 2024; but see Rossi et al., 2021; Tourmente et al., 2011; Quintero-Pérez et al., 2023).

Sperm are among the most evolutionarily labile cell types, with extensive diversification in morphology across taxa (Pitnick et al., 2009; Lüpold and Pitnick, 2018; Afzelius, 1995; Kahrl et al., 2022). Whether or not this lability extends to sperm cell thermal traits is not well understood, though there is some evidence that it does (Rahman et al., 2009; Tourmente et al., 2011; Lahnsteiner and Mansour, 2012; Poullet et al., 2015; Revathy and Benno Pereira, 2016; Andronikov, 1975). If sperm heat tolerance can evolve within a population, one would expect repeatable variation in sperm heat tolerance among males. Repeatability sets an upper bound on heritability in a given environmental context, and is often positively correlated with observed heritability (Boake, 1989; Dohm, 2002; Morgan et al., 2018; Bell et al., 2009). The repeatability of sperm heat tolerance is completely unknown.

One might also expect co-adaptation of sperm and adult thermal tolerance, since adult performance in a thermal environment is moot if reproduction is compromised (Angilletta et al. 2006). Consistent with this prediction, though not about sperm cells *per se*, adult and fertility heat tolerance are correlated across species of *Drosophila* (van Heerwaarden and Sgrò 2021; Wang and Gunderson 2022). Thermal co-adaptation is also expected to occur at the intraspecific level (Angilletta et al. 2006; David et al., 2005). However, to our knowledge this has not been tested with respect to sperm thermal tolerance in animals.

One limitation to measuring sperm cell thermal traits is that isolating and keeping sperm alive long enough for experimentation is challenging, and in some cases impossible. In many species, sperm cells either do not survive or perform poorly outside the body (Pitnick, 2009; Browne et al., 2015; Pennington, 1985; Orr and Brennan, 2015). In addition, the lifespan of ejaculated sperm in many taxa is extremely short, making data collection difficult. For example, the sperm of many external fertilizers die within seconds of ejaculation (Alavi and Cosson, 2005; Browne et al., 2015). Finally, in some taxa sperm must be collected destructively, making estimates of within-individual repeatability untenable.

Here, we used the Cuban brown anole lizard (*Anolis sagrei*) as a model organism to investigate sperm heat sensitivity. Anoles are an excellent system for our purposes because their sperm are easy to collect, are relatively long-lived *in vitro*, and can be sampled non-destructively, permitting repeated measurement of sperm traits on the same individual (Kahrl and Cox, 2015, 2017). Using a novel approach we developed to measure sperm cell heat tolerance, we address three questions. First, is sperm heat tolerance repeatable at the individual level, indicative of evolutionary potential? Second, is there a correlation between sperm and adult heat tolerance, consistent with thermal co-adaptation? Third, does episodic heat stress experienced by males compromise sperm performance, consistent with high heat sensitivity of reproductive potential?

## Methods

### Lizard collection and housing

N = 31 adult male *A. sagrei* were collected from New Orleans, Louisiana, USA (29.9407° N, 90.1203° W) in June and July, within their reproductive season (Kahrl and Cox, 2015; Tokarz, 1998). Lizards were immediately transported to Tulane University where they were housed individually in plastic cages (18×11×14 cm) in a climate-controlled growth chamber (Percival, I30NLC9) set to a thermal regime that mimics mean summer temperatures in New Orleans (24-35°C night:day fluctuating temperatures, 75 % relative humidity, 12h:12h light:dark cycle; Deery et al. 2021). Lizards were watered daily and fed crickets dusted with calcium and vitamin powder three times per week. All animal protocols were approved by the Tulane University Institutional Animal Care and Use Committee (protocol #1436).

### Sperm extraction and measurement of heat tolerance

The heat tolerance of sperm from each lizard was measured four times over 21 days (Fig. 1). The initial Day 0 sample was collected and measured on the day of capture, and the rest were collected every seven days. Semen was extracted by gently pressing on the pelvic girdle, causing ejaculate to emerge from the cloaca (Kahrl and Cox, 2015; Kahrl et al., 2019). Approximately 2μL of semen was collected with a pipette and immediately diluted in 1000ul Eagle’s minimal essential medium (DMEM) in a microcentrifuge tube (Kahrl and Cox, 2015; Kahrl et al., 2019). Sperm heat tolerance was taken as the temperature at which 50% of sperm from an ejaculate sample become immobile (LT50) after 30 min of exposure. Each diluted ejaculate sample was split into seven 40 μL sub-samples in a 96-well PCR plate. Each sub-sample was exposed to one of the following temperatures: 33, 36, 38, 41, 43, 46 or 50°C using the gradient function of a PCR thermocycler (Eppendorf, Mastercycler® X50). After thermal exposure, 10 μL of each sample was removed with a pipette and loaded into a hemocytometer. Samples were video recorded for 20 seconds under a compound light microscope (Nikon Eclipse) connected to a digital camera (Nikon D3s). Videos were viewed with the imageJ plugin “cell counter,” and the percentage of sperm that were motile was recorded. To calculate the LT50 of an ejaculate, we first normalized the motility of each sub-sample by dividing every motility value by the maximum value across all seven sub-samples. This normalization accounts for variation in baseline sperm motility among different lizards and removes the effect of baseline sperm motility on LT50 calculation (Motulsky, 1999; Weimer et al., 2012). LT50 was calculated by fitting a dose-response model to the data for a given ejaculate using a four-parameter log-logistic model with the LL.4 function in the R package ‘drc’ (Ritz and Streibig 2005; Ritz et al. 2015; Parratt et al., 2021; van Heerwaarden and Sgrò 2021). We defined “baseline” sperm motility as the motility of the subsample incubated at 33°C because this temperature is within the preferred body temperature range of *A. sagrei* (Ryan and Gunderson 2021).

**Fig. 1.**
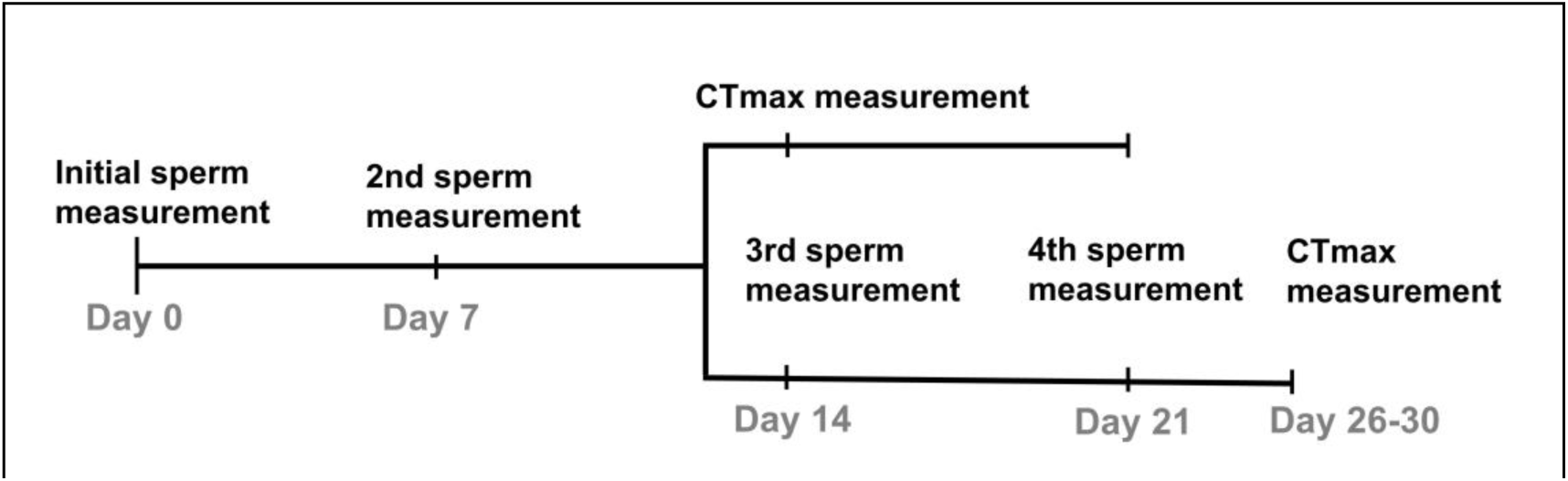
Overview of the experimental design. Field-collected adult males were brought to the laboratory for sperm and adult trait measurement over several weeks. Initial sperm traits (baseline sperm motility and heat tolerance) were measured on the day of capture (Day 0), after which sperm heat tolerance was measured every seven days for three weeks. On Day 14, adult heat tolerance (CTmax) was measured for 15 of the individuals prior to sperm heat tolerance measurement. For the remaining lizards, CTmax was measured at the end of the experiment (Day26 to 30), after the Day 21 sperm samples were collected.

### Adult heat tolerance

Adult heat tolerance was measured as the Critical Thermal Maximum (CTmax), the temperature at which a lizard loses neuromuscular coordination and cannot right itself during a thermal ramp. We followed published protocols for anoles (Leal and Gunderson, 2012; Gunderson et al., 2018; Deery et al., 2021). Lizards were loosely tethered inside a cardboard enclosure (30×30×16cm) with dental floss around the waist and placed under a 100W incandescent lightbulb. To monitor body temperature in real time, the tip of a T-type 36-gauge wire thermocouple probe connected to a digital thermometer (Omega HH81A) was inserted into the cloaca and secured with a small amount of super glue. Body heating rate was maintained at 2°C/min for all animals (Gunderson, 2024). Lizards were flipped on their backs to test the righting response at 1°C body temperature intervals starting at 38°C. If the lizard righted itself within 10 sec, it was returned under the heat lamp and flipped again at the next test temperature. If the lizard was unable to right itself the trial ended. CTmax was recorded as 0.5 °C above the highest body temperature at which the animal righted itself (Deery et al., 2021).

### Testis size measurement

At the conclusion of the experiment, lizards were anaesthetized with isoflurane and euthanized via cervical dislocation. Testis were removed, patted dry with a paper towel, and wet weight was measured with a digital scale (VWR-64B2T).

### Data analysis

We tested for repeatability of sperm heat tolerance and baseline motility in two ways. First, we tested for a correlation between measurements from Day 0 and Day 7 ejaculate samples with linear regression using the ‘lm’ function in R (R core team, 2022). Data from all 31 animals were included in these analyses. Second, we calculated the intraclass correlation coefficient (ICC) across all samples with mixed-effects models with individual identity as a random factor using the ‘lme4 ‘package in R (Bates et al. 2015; Morgan et al., 2018; Wolak et al., 2012; Lessells and Boag, 1987). Only animals that did not experience CTmax measurement on Day 14 (N = 16; Fig. 1) were included to remove confounding effects of different thermal experience among individuals.

To test for an association between adult and sperm heat tolerance, we conducted linear regression using the ‘lm’ function in R (R core team, 2022). The model included sperm LT50 as the response variable with CTmax and the day of CTmax measurement as explanatory variables. Sperm LT50 measurements from Day 7 were used for individuals that had CTmax measured on Day 14, and sperm LT50 measurements from Day 21 were used for those individuals that had CTmax measured at the end of the experiment.

To test for relationships between sperm traits, body condition, and testis size, we conducted multiple linear regression using the ‘lm’ function in R, with gonadosomatic index (GSI) and body mass index (BMI) as predictors, and baseline sperm motility or sperm LT50 on Day 7 as response variables. GSI was calculated by dividing the wet mass of the gonads by lizard body mass (Noeske and Meier, 1977; Pearson et al., 1976; Tourmente et al., 2011). BMI, a measure commonly used for body condition (eg., Battles et al. 2013, Vervust et al. 2008; Goodman 2008), was obtained by dividing individual mass by the square of snout-vent length (SVL). We chose the Day-7 measurement, following a week of acclimation in the lab, to minimize the effects of individual variation induced by prior experience in the field.

To test for an effect of adult heat exposure on sperm, we used paired t-tests to model changes in sperm LT50 and baseline sperm motility between Day 7 (before CTmax measurement) and Day 14 (after CTmax measurement) ejaculate samples for individuals that did and did not experience CTmax trials (Fig. 1).

## Results

### Sperm heat tolerance

Across individuals, sperm motility was similar after exposure to temperatures from 33-41°C, but began to notably decrease at higher temperatures (Fig. 2). The initial (Day 0) sperm LT50 varied among individuals (range 42.2-45.2°C), with a mean (±SE) of 43.8±0.1°C (Fig. 2). Initial baseline sperm motility (percent motile sperm after 33°C incubation) also varied considerably among males (range 59 - 98 %; Fig. 2), with a mean of 86.1 ± 1.9%.

**Fig. 2.**
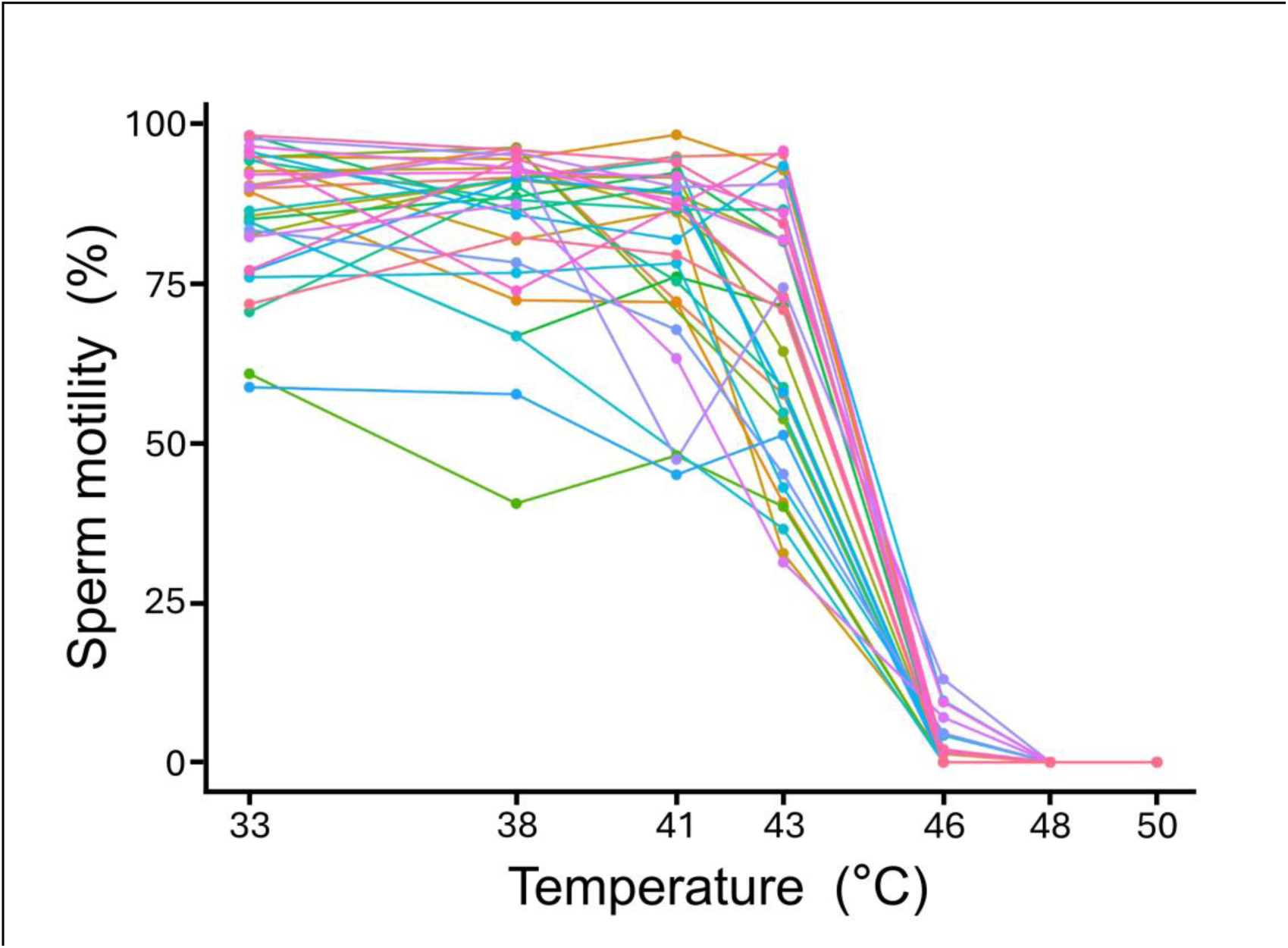
Percentage of sperm motile after 30 min exposed to different temperatures. Examples here are from samples collected on Day 0. Lines connect sperm derived from the same ejaculate of a single individual (N=31).

### Sperm trait repeatability

There was considerable repeatability in sperm heat tolerance. Among all 31 individuals, there was a significant positive correlation between Day 0 and Day 7 sperm LT50 (Fig. 3A; R-sq = 0.22, F=8.34, *P* = 0.007). Among the 16 individuals that did not undergo CTmax measurement on Day 14 (Fig. 1), the repeatability of sperm LT50 across the four ejaculate samples was 0.17 (Fig. 3B). We also calculated repeatability with just the Day 7 through Day 21 samples such that all samples were collected after males had been acclimated to laboratory conditions. Repeatability was considerably higher in this analysis (0.38). Additionally, sperm LT50 significantly increased over the course of the experiment, from 43.8 ± 0.1 °C in the initial Day 0 sample to 44.5 ± 0.1 °C in the Day 21 sample (Fig.3B, linear mixed-effects model: F = 4.25, *P* < 0.001).

**Fig. 3.**
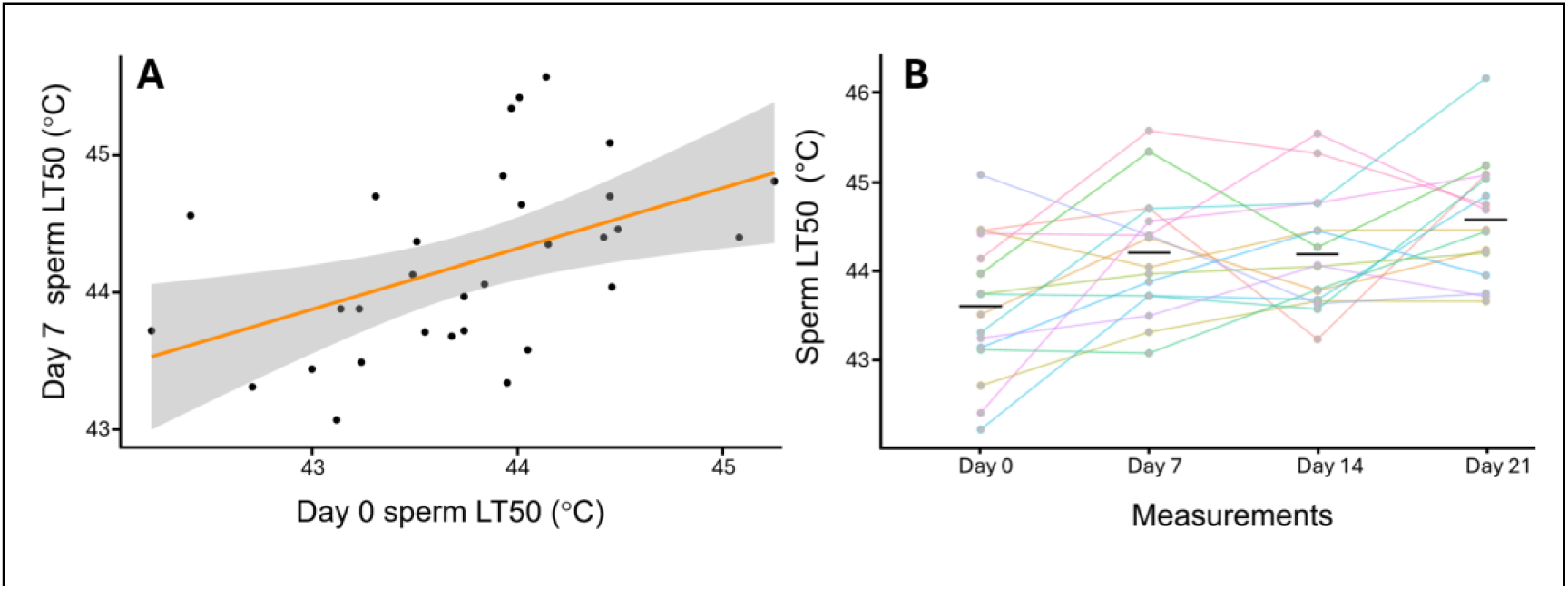
Repeatability in sperm heat tolerance. **A)** Correlation between sperm heat tolerance (LT50) from the same male on Day 0 versus Day 7 (*P* = 0.007). The gray area denotes the 95% confidence interval of the best-fit line. **B)** Sperm heat tolerance across the study for animals that did not experience CTmax measurement on Day 14. Each line represents one individual male. The black bars indicate the mean for each day.

Baseline sperm motility (motility after incubation at 33°C) was also repeatable. There was a significant positive correlation between baseline sperm motility measured on Day 0 and Day 7 (Fig. 4A, R-sq = 0.15, *P* = 0.033). The overall repeatability of baseline sperm motility across 4 trials was 0.51 for the 16 lizards that did not undergo CTmax trials on Day 14 (Fig. 4B). Repeatability was similar (0.52) considering only the data collected on Days 7-21, after laboratory acclimation.

**Fig. 4.**
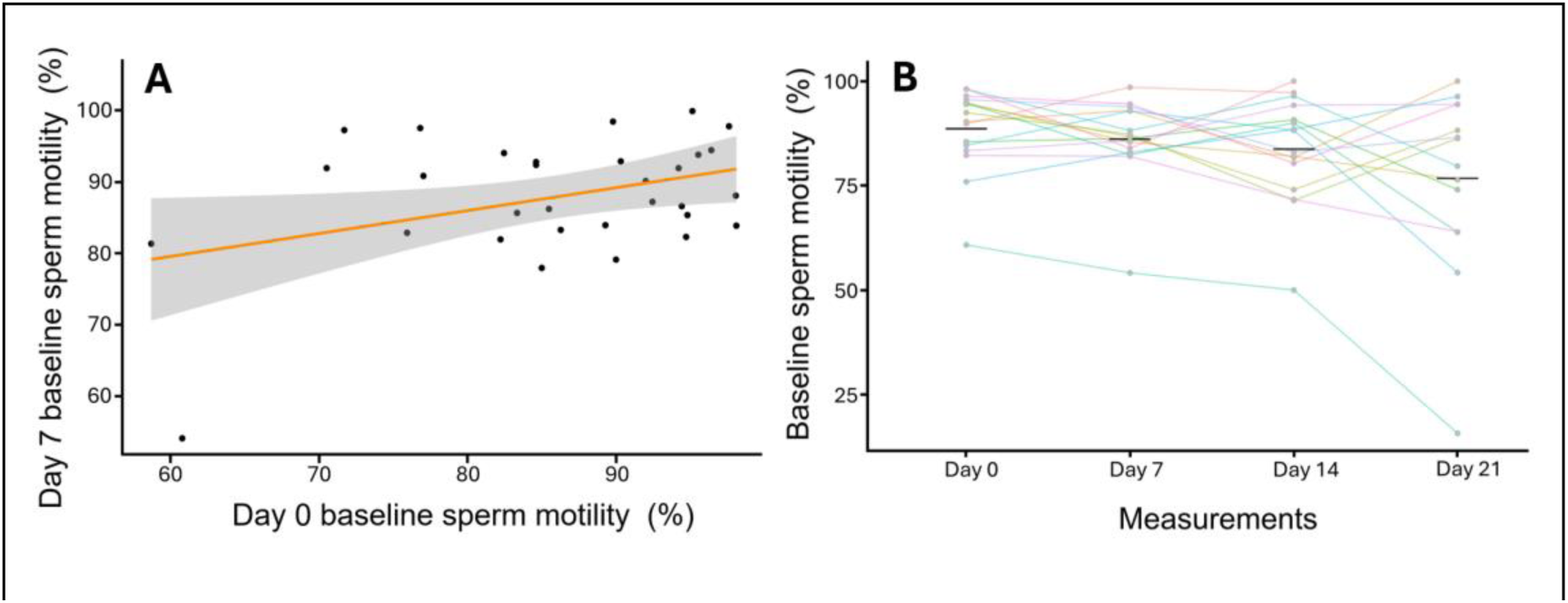
Repeatability in baseline sperm motility. **A)** Correlation between baseline sperm motility on Day 0 versus Day 7 (*P* = 0.033). The grey area denotes the 95% confidence interval of the best-fit line. **B)** Baseline sperm motility across the study for animals that did not experience CTmax measurement on Day 14. Each line represents one individual male. The black bars indicate the mean for that day.

### Associations between adult and sperm traits

There was no correlation between adult and sperm heat tolerance (Fig. 5; N=28, *P*= 0.621). Furthermore, neither adult body condition (BMI) nor gonadosomatic index (GSI) were associated with sperm heat tolerance or baseline motility (Sperm LT50: GSI N=28, *P* = 0.701, BMI N=28, *P* = 0.142; Baseline motility: GSI N=28, *P* = 0.828, BMI N = 28, *P* = 0.981; Table 1).

**Fig. 5.**
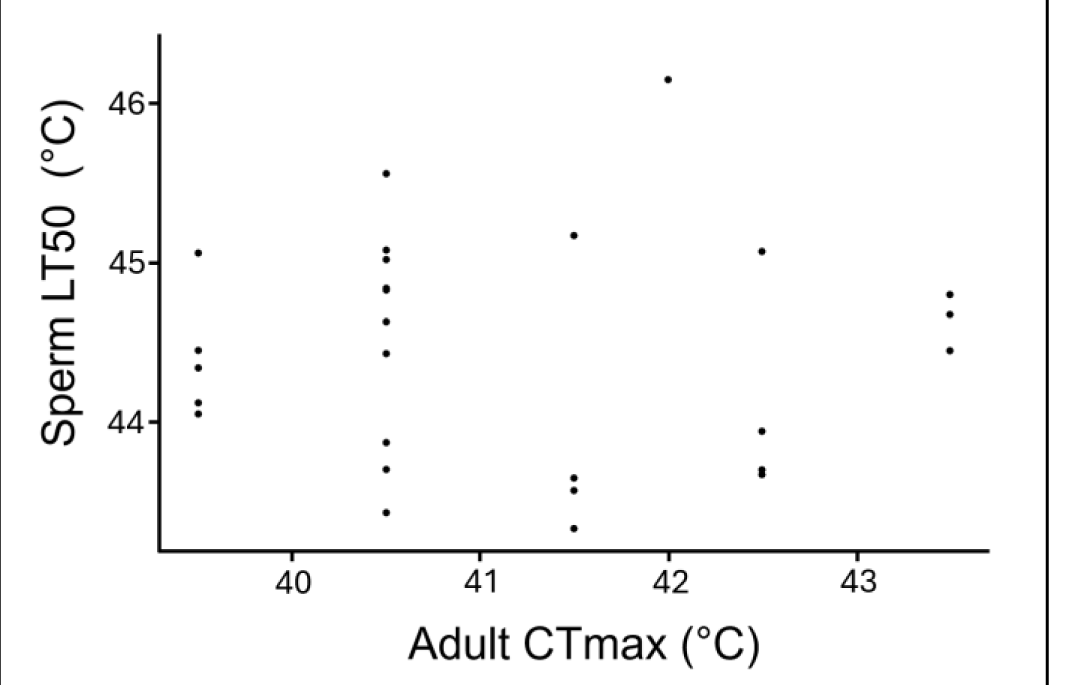
No association between adult and sperm heat tolerance. Adult CTmax and sperm LT50 were unrelated (*P* = 0.621).

**Table 1.** Associations between adult and sperm traits. Results from multiple linear regression models testing for relationships between sperm traits, body condition (BMI), and testis size (GSI).

| Trait | Predictors | Std. error | t-value | p-value |
| --- | --- | --- | --- | --- |
| **Baseline motility** | Intercept | 14.207 | 6.003 | <0.001 |
|  | GSI | 3.969 | 0.219 | 0.828 |
|  | BMI | 1.143 | 0.023 | 0.981 |
| **Sperm LT50** | Intercept | 0.983 | 46.572 | <0.001 |
|  | GSI | 0.274 | -0.388 | 0.701 |
|  | BMI | 0.079 | -1.520 | 0.142 |

### Adult heat exposure and sperm performance

To evaluate if episodic male heat stress impacts sperm, we compared sperm traits of lizards that did and did not undergo CTmax measurement (Fig. 1). Baseline sperm motility decreased significantly for the group exposed to their CTmax (mean change = -11.2 ± 3.6; paired t-test: t(14)=2.991, *p* = 0.009) but not in the non-exposed group (- 2.4 ± 2.1; paired t-test: t(15)=0.972, *P* = 0.346; Fig. 6A). Sperm heat tolerance did not change in the CTmax exposed (t(14)=0.383, *P* = 0.707) or non-exposed group (t(15)=0.278, *P* = 0.784; Fig. 6B).

**Fig. 6.**
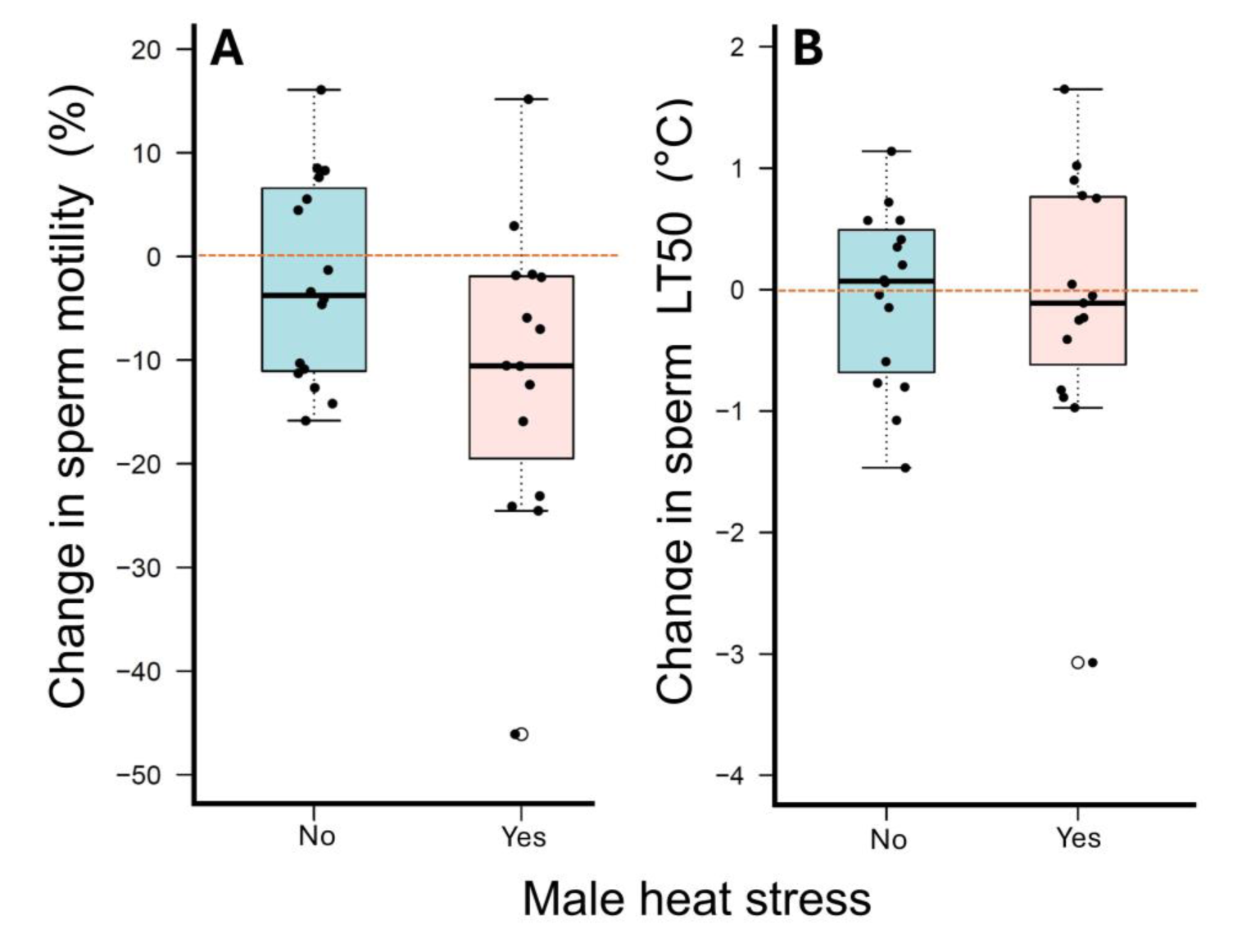
Adult heat exposure and sperm performance. **A)** Effect of adult heat exposure on baseline sperm motility. The change is calculated as the difference in baseline motility between Day 14 (post-CTmax heat exposure) and Day 7 (pre-CTMax). **B)** Effect of adult heat exposure on sperm LT50.

## Discussion

We aimed to understand how gametes respond to heat in a lizard model using a novel approach to test the heat tolerance of ejaculated sperm cells. We found repeatable individual-level variation in sperm heat tolerance and baseline motility, a lack of association between adult and sperm heat tolerance, and an effect of a single, brief extreme heat exposure on sperm performance. We discuss these results in turn below.

### Repeatability in Sperm Traits

There was considerable consistency in sperm heat tolerance among males. First, there was a positive correlation between sperm heat tolerance measurements taken on different days (Fig. 3A). Second, sperm heat tolerance repeatability was 0.17 across the entire experiment (Days 0-21) and 0.38 when restricted to samples collected after lab acclimation (Days 7-21; Fig. 3B). These results suggest consistent individual-level variation in sperm heat tolerance within the population. We know of no other estimates of repeatability in sperm heat tolerance, but calculation of adult heat tolerance repeatability yields similar values in other vertebrates (Morgan et al., 2018; Grinder et al., 2020), ranging from 0.1 in the grey-cheecked salamander (*Plethodon metacalfi*; McTernan and Sears, 2022) to 0.48 in brook trout (*Salvelinus fontinalis*; O’Donnell et al. 2020). The repeatability of baseline sperm motility (0.52; Fig. 4) was even greater, and higher than observations in other vertebrates. For example, sperm motility was not repeatable (<0.01) in house sparrows (*Passer domesticus*) and Spanish sparrows (*P. hispanio lensis*; Cramer et al., 2015). The repeatability of sperm viability was 0.18 in rabbits (*Oryctolagus cuniculus;* García-Tomás et al., 2006) and 0.35 in rams (*Ovis aries;* Langford et al., 1989).

Repeatability alone does not demonstrate that trait variation is genetic, but it does set the upper limit for, and is commonly correlated with, heritability (Boake, 1989; Dohm, 2002; Morgan et al., 2018; Bell et al., 2009). The repeatability of sperm heat tolerance and baseline motility in brown anoles is therefore consistent with evolutionary potential in these traits. That said, our data also provide clear evidence that environmental conditions influence sperm traits. For example, sperm heat tolerance increased as lizards spent more time in the laboratory (Fig 3 B). In addition, the repeatability of sperm heat tolerance more than doubled after animals were lab-acclimated. These finding emphasizes the intricacies of predicting evolutionary responses to climate change given the interaction between environmental and genetic factors in influencing the expression of physiological traits (Dowd et al., 2015; Jimenez et al., 2015).

Sperm traits are sometimes linked to individual body condition, perhaps reflecting variation in stress and/or resource acquisition prior to capture (Gage and Cook, 1994; Kahrl and Cox, 2015; Malo et al., 2005; Rahman et al., 2013). For example, sperm number, morphology, and fertilization success under sperm competition were correlated with male body condition in a different population of brown anoles (Kahrl and Cox, 2015). In some cases, this correlation extends to testes size, which can be indicative of reproductive investment. For instance, in red deer (*Cervus elaphus*), sperm velocity was associated with relative testes size. However, we found no evidence for condition-dependent sperm performance in our population. Neither body condition (BMI) or relative testis size (GSI) correlated with sperm heat tolerance or baseline motility (Table 1). Why some sperm traits associate with condition and others do not is unclear, but may reflect differences in the relative cost of maintaining them (Kahrl and Cox, 2015).

### Sperm and adult heat tolerance are not correlated

Although data are sparse, interspecific studies often find that species with heat tolerant adults produce more heat tolerant sperm (Rahman et al., 2009; Tourmente et al., 2011; Lahnsteiner and Mansour, 2012; Poullet et al., 2015; Revathy and Benno Pereira, 2016; reviewed in Wang and Gunderson 2022). This pattern is consistent with the expected co-adaptation of different thermal traits (Angilletta et al. 2006) and genetic linkage between adult and gamete thermal tolerance at a macroevolutionary scale. Whether this holds at the intraspecific level is unknown. If sperm and adult thermal tolerance are genetically linked, it would suggest that these traits will easily co-evolve, and that thermal adaptation can potentially happen more quickly because of temperature-driven selection at the gamete level (Clarke et al., 2004; Mohapatra et al., 2020). However, we found no association between adult and sperm thermal tolerance in brown anoles at the individual level (Fig. 5). More data directly comparing sperm and adult tolerance would be valuable to understand linkages between adult and sperm thermal tolerance across intra- and inter-specific scales.

### Episodic adult heat stress decreases sperm performance

With global climate change, organisms are more likely to encounter extreme heat events (Christidis et al., 2015; Meehl and Tebaldi, 2004; Sales et al., 2018) that can potentially threaten biodiversity through reduced fertility (David et al., 2005; Sales et al., 2018, 2021; Vasudeva et al., 2021; Walsh et al., 2019, 2022; McAfee et al., 2020; Rinehart et al., 2000). We found that males exposed to a single, brief heat extreme (CTmax experiments last less than 10 min, and because heat is ramped, the maximum temperature is sustained for less than 20 sec) had decreased sperm motility (Fig. 6). The observed effect must have been on stored rather than developing sperm because semen was collected immediately after heat exposure and spermatogenesis takes weeks (Sharma and Agarwal, 2011; Gribbins, 2011). Decreased motility in stored sperm cells could contribute to observed heat effects in other system in which sperm were not directly measured. For example, in many *Drosophila* species, males temporarily lose fertility after heat exposure (David, 2005; Jørgensen et al., 2006; Parratt et al., 2021; Sutter et al., 2019). Our findings also align with recent studies on flour beetles (*T. castaneum*), where pre-mating heat exposure impacted sperm traits and fertility (Sales et al., 2018, 2021; Vasudeva et al., 2021). Significant impacts of heat on male fertility appear to be a widespread across animal taxa, from insects to reptiles, raising concern about the vulnerability of reproduction to warming (Walsh et al., 2019; Wang and Gunderson 2022).

## Conclusion

Organisms must maintain reproductive capability to persist under rapid global climate change. However, there is increasing evidence that reproduction is particularly vulnerable to warming (Wang and Gunderson, 2022; Sales et al., 2021; Walsh et al., 2019, 2022). We provide evidence for heat sensitivity in lizard sperm cells and environmental effects on sperm performance more generally, but also considerable repeatability in sperm thermal traits consistent with evolutionary potential. Our work reinforces that lizards are an excellent model system for investigating sperm cell thermal biology, given that their sperm can be extracted repeatedly and non-destructively and have relatively long lifespans outside of the body (Rossi et al., 2021; Kahrl and Cox, 2015, 2017). Overall, this study provides important context for future investigations into the resilience and adaptability of reproductive traits in the increasingly challenging thermal environments created by climate change.

### Data accessibility

Data are available in a GitHub repository: https://github.com/wwywanglizardsperm/sperm_thermal_data.git

## Acknowledgements

We would like to thank all members of the Gunderson Lab who helped with animal care and engaging in helpful discussions about this project, additionally Jalen LaCour and Jess Valentine for helping with data collection. We thank Dr. Kyriakos Papadopoulos and Chia-Yu Chen for helping with preliminary data collection. We also thank Scott Pitnick, Hung-Lun Tu and Rou-Rung Chen for helpful discussions and feedback, and Kuan-Jui Su for his kind assistance with the R script.

## References

1. Afzelius, B. A. (1995). Gustaf Retzius and spermatology. Int. J. Dev. Biol. 39, 675–685

2. Alavi, S. M. and Cosson, J. (2005). Sperm motility in fishes. I. Effects of temperature and pH: a review. Cell Biol. Internat. 29, 101–110. doi: 10.1016/j.cellbi.2004.11.021

3. Andronikov, V. B. (1975). Heat resistance of gametes of marine invertebrates in relation to temperature conditions under which the species exist. Mar. Biol. 30, 1–11

4. Bates, D., Mächler, M., Bolker, B. and Walker, S. (2015). Fitting Linear Mixed-Effects Models Using lme4. J. Stat. Softw. 67, 1–48. 10.18637/jss.v067.i01

5. Battles, A. C., Whittle, T. K., Stehle, C. M. and Johnson, M. A. (2013). Effects of human land use on prey availability and body condition in the green anole lizard, Anolis carolinensis. Herpetol. Conserv. Bio. 8, 16–26.

6. Binet, M. T. and Doyle C. J. (2013). Effect of near-future seawater temperature rises on sea urchin sperm longevity. Mar. Freshwater Res. 64, 1–9. 10.1071/Mf12121

7. Browne, R. K., Kaurova, S. A., Uteshev, V. K., Shishova, N. V., McGinnity, D., Figiel, C. R., Mansour, N., Agney, D., Wu, M., Gakhova, E. N., Dzyuba, B. and Cosson, J. (2015). Sperm motility of externally fertilizing fish and amphibians. Theriogenology 83, 1–13. 10.1016/j.theriogenology.2014.09.018

8. Christidis, N., Jones, G. S. and Stott, P. A. (2015). Dramatically increasing chance of extremely hot summers since the 2003 European heatwave. *Nat*. Clim. Change 5, 46–50. 10.1038/nclimate2468

9. Cramer, E. R., Laskemoen, T., Stensrud, E., Rowe, M., Haas, F., Lifjeld, J. T., Saetre, G. P. and Johnsen, A. (2015). Morphology-function relationships and repeatability in the sperm of Passer sparrows. J. Morphol. 276, 370–377. 10.1002/jmor.20346

10. David, J. R., Araripe, L. O., Chakir, M., Legout, H., Lemos, B., Pétavy, G., Rohmer, C., Joly, D. and Moreteau, B. (2005). Male sterility at extreme temperatures: a significant but neglected phenomenon for understanding *Drosophila* climatic adaptations. J. Evol. Biol. 18, 838–846. 10.1111/j.1420-9101.2005.00914.x

11. Deery, S. W., Rej, J. E., Haro, D. and Gunderson, A. R. (2021). Heat hardening in a pair of *Anolis* lizards: constraints, dynamics and ecological consequences. J. Exp. Biol. 224, jeb240994. 10.1242/jeb.240994

12. Dougherty, L. R., Frost, F., Maenpaa, M. I., Rowe, M., Cole, B. J., Vasudeva, R., Pottier, P., Schultner, E., Macartney, E. L., Lindenbaum, I. and Smith, J. L., et al. (2024). A systematic map of studies testing the relationship between temperature and animal reproduction. Ecol. Solut. Evid. 5, e12303. 10.1002/2688-8319.12303

13. Dowd, W. W., King, F. A. and Denny, M. W. (2015). Thermal variation, thermal extremes and the physiological performance of individuals. J. Exp. Biol, 218, 1956–1967. 10.1242/jeb.114926

14. Gage, M. J. G. and Cook, P. A. (1994). Sperm size or numbers? Effects of nutritional stress upon eupyrene and apyrene sperm production strategies in the moth *Plodia interpunctella* (Lepidoptera: Pyralidea). Funct. Ecol. 8, 594. 10.2307/2389920

15. Goodman R. M. (2008). Latent effects of egg incubation temperature on growth in the lizard *Anolis carolinensis*. J. Exp. Zool. 309, 525–533. 10.1002/jez.483

16. Gribbins, K. (2011). Reptilian spermatogenesis: A histological and ultrastructural perspective. Spermatogenesis 1, 250–269. 10.4161/spmg.1.3.18092

17. Grinder, R. M., Bassar, R. D. and Auer, S. K. (2020). Upper thermal limits are repeatable in Trinidadian guppies. J. Therma. Biol. 90, 102597. 10.1002/jez.483doi.org/10.1016/j.jtherbio.2020.102597

18. Gunderson, A. R. (2024). Disentangling physiological and physical explanations for body size-dependent thermal tolerance. J. Exp. Biol. 227, doi: 10.1242/jeb.245645.

19. Huey, R. B., Kearney, M. R., Krockenberger, A., Holtum, J. A., Jess, M. and Williams, S. E. (2012). Predicting organismal vulnerability to climate warming: roles of behavior, physiology and adaptation. Phil. Trans. R. Soc. B, 367, 1665–1679. 10.1098/rstb.2012.0005

20. Hughes, L, (2000). Biological consequences of global warming: is the signal already apparent? Trends Ecol. Evol. 15, 56–61. 10.1016/s0169-5347(99)01764-4

21. IPCC (2014). “Climate Change 2014: Synthesis report,” in contribution of working groups I, II and III to the fifth assessment report of the intergovernmental panel on climate change, eds R. K. Pachauri and L. A. Meyer (Geneva: IPCC), 151.

22. Jimenez, A. G., Jayawardene, S., Alves, S., Dallmer, J. and Dowd, W. W. (2015). Micro-scale environmental variation amplifies physiological variation among individual mussels. Proc. Biol. Sci. 282, 20152273. 10.1098/rspb.2015.2273

23. Jørgensen, K. T., Sørensen, J. G. and Bundgaard, J. (2006). Heat tolerance and the effect of mild heat stress on reproductive characters in *Drosophila buzzatii* males. J. Therm. Biol. 31, 280–286. 10.1016/j.jtherbio.2005.11.026

24. Kahrl, A. F. and Cox, R. M. (2017). Consistent differences in sperm morphology and testis size between native and introduced populations of three *Anolis* lizard species. J. Herpetol. 51, 532–537. 10.1670/16-184

25. Kahrl, A. F. and Cox, R. M. (2015). Diet affects ejaculate traits in a lizard with condition-dependent fertilization success. Behav. Ecol. 26, 1502–1511. 10.1093/beheco/arv105

26. Kahrl, A. F. Johnson, M. A., and Cox, R. M. (2019). Rapid evolution of testis size relative to sperm morphology suggests that post-copulatory selection targets sperm number in *Anolis* lizards. J. Evol. Biol. 32, 302–309. 10.1111/jeb.13414

27. Lahnsteiner, F. and Mansour, N. (2012). The effect of temperature on sperm motility and enzymatic activity in brown trout Salmo trutta, burbot *Lota lota* and grayling *Thymallus thymallus*. J. Fish Biol. 81, 197–209. doi:10.1111/j.1095-8649.2012.03323.x

28. Langford, G. A., Shrestha, J. N. B. and Marcus, G. J. (1989). Repeatability of scrotal size and semen quality measurements in rams in a short-day light regime. Anim. Reprod. 19, 19–27. 10.1016/0378-4320(89)90043-2

29. Lessells, C. M. and Boag, P. T. (1987). Unrepeatable repeatabilities: A common mistake. The Auk, 104, 116–121. 10.2307/4087240

30. Lüpold, S. and Pitnick, S. (2018). Sperm form and function: what do we know about the role of sexual selection? Reproduction, 155, R229–R243. 10.1530/REP-17-0536

31. Malo, A. F., Roldan, E. R. S., Garde, J., Soler, A. J. and Gomendio, M. (2005). Antlers honestly advertise sperm production and quality. Proc. Biol. Sci. 272, 149–157. 10.1098/rspb.2004.2933

32. McAfee, A., Chapman, A., Higo, H., Underwood, R., Milone, J., Foster, L. J., Guarna, M. M., Tarpy, D. R. and Pettis, J. S. (2020). Vulnerability of honey bee queens to heat-induced loss of fertility. Nat. Sustain. 3, 367–376. 10.1038/s41893-020-0493-x

33. Meehl, G. A. and Tebaldi, C. (2004). More intense, more frequent, and longer lasting heat waves in the 21st century. Science, 305, 994–997. 10.1126/science.1098704

34. Morgan, R., Finnøen, M. H. and Jutfelt, F. (2018). CTmax is repeatable and doesn’t reduce growth in zebrafish. Sci. Rep. 8, 1–8. 10.1038/s41598-018-25593-4

35. Motulsky, H. and Christopoulos, A., 2004. Fitting Models to Biological Data Using Linear and Nonlinear Regression. A Practical Guide to Curve Fitting. Oxford University Press, New York, NY.

36. Noeske, T. A. and Meier, A. H. (1977). Photoperiodic and thermoperiodic interaction affecting fat stores and reproductive indexes in the male green anole, *Anolis carolinensis*. J. Exp. Zool. 202, 97–102. 10.1002/jez.1402020112

37. Orr, T. J. and Brennan, P. L. (2015). Sperm storage: distinguishing selective processes and evaluating criteria. Trends Ecol. Evol. 30, 261–272. 10.1016/j.tree.2015.03.006.

38. Parratt, S. R., Walsh, B. S., Metelmann, S., White, N., Manser, A., Bretman, A. J., Hoffmann, A. A., Snook, R. R. and Price, T. A. R. (2021). Temperatures that sterilize males better match global species distributions than lethal temperatures. *Nat*. Clim. Change 11, 481–484. 10.1038/s41558-021-01047-0

39. Pearson, A. K., Tsui, H. W. and Licht, P. (1976). Effect of temperature on spermatogenesis, on the production and action of androgens and on the ultrastructure of gonadotropic cells in the lizard *Anolis carolinen*sis. J. Exp. Zool, 195, 291–303. 10.1002/jez.1401950214

40. Pitnick, S., Hosken, D. J. and Birkhead, T. R. (2009). Sperm Biology: An Evolutionary Perspective. *San Diego*: Academic Press.

41. Poullet, N., Vielle, A., Gimond, C., Ferrari, C. and Braendle, C. (2015). Evolutionarily divergent thermal sensitivity of germline development and fertility in hermaphroditic *Caenorhabditis* nematodes. Evol. Dev. 17, 380–397. doi: 10.1111/ede.12170

42. R Core Team (2022). R: A language and environment for statistical computing. R Foundation for Statistical Computing, Vienna, Austria. URL https://www.R-project.org/.

43. Rahman, M. M., Kelley, J. L. and Evans, J. P. (2013). Condition-dependent expression of pre- and postcopulatory sexual traits in guppies. Ecol. Evol. 3, 2197–2213. 10.1002/ece3.632

44. Rahman, M. S., Tsuchiya, M. and Uehara, T. (2009). Effects of temperature on gamete longevity and fertilization success in two sea urchin species, *Echinometra mathaei* and *Tripneustes gratilla*. Zoolog. Sci. 26, 1–8. doi: 10.2108/zsj.26.1

45. Revathy, S. and Benno Pereira, F. G. (2016). Effect of temperature on sperm motility in fishes. Internat. J. Sci. Res. 5, 81–89.

46. Rinehart, J. P., Yocum, G. D. and Denlinger, D. L. (2000). Thermotolerance and rapid cold hardening ameliorate the negative effects of brief exposures to high or low temperatures on fecundity in the flesh fly, *Sarcophaga crassipalpis*. Physiol. Entomol. 25, 330–336. doi: 10.1111/j.1365-3032.2000.00201.x

47. Ritz, C., Baty F, Streibig J. C. and Gerhard, D (2015). Dose-response analysis using R. PLoS ONE 10, e0146021. 10.1371/journal.pone.0146021

48. Ritz, C. and Streibig, J. C. (2005). Bioassay Analysis Using R. J. Stat. Softw. 12, 1–22. 10.18637/jss.v012.i05

49. Rohmer, C., David, J. R., Moreteau, B. and Joly, D. (2004). Heat induced male sterility in *Drosophila melanogaster*: adaptive genetic variations among geographic populations and role of the Y chromosome. J. Exp. Biol. 207, 2735–2743. 10.1242/jeb.01087

50. Rossi, N., Lopez Juri, G., Chiaraviglio, M. and Cardozo, G. (2021). Oviductal fluid counterbalances the negative effect of high temperature on sperm in an ectotherm model. Biol. Open 10, 058593. doi: 10.1242/bio.058593

51. Sales, K., Vasudeva, R. and Gage, M. J. G. (2021). Fertility and mortality impacts of thermal stress from experimental heatwaves on different life stages and their recovery in a model insect. R. Soc. Open Sci., 8, 201717. 10.1098/rsos.201717

52. Sales, K., Vasudeva, R., Dickinson, M. E., Godwin, J. L., Lumley, A. J., Michalczyk, Ł., Hebberecht, L., Thomas, P., Franco, A. and Gage, M. J. G. (2018). Experimental heatwaves compromise sperm function and cause transgenerational damage in a model insect. Nat. Commun. 9, 4771. 10.1038/s41467-018-07273-z

53. Sharma, R. and Agarwal, A. (2011). Spermatogenesis: An overview. Sperm Chromatin. Berlin, Germany: Springer.

54. Sutter, A., Travers, L. M., Oku, K., Delaney, K. L., Store, S. J., Price, T. A. R. and Wedell, N. (2019). Flexible polyandry in female flies is an adaptive response to infertile males. Behav. Ecol. 30, 1715–1724. 10.1093/beheco/arz140

55. Tokarz, R. R. (1998). Mating pattern in the lizard *Anolis sagrei*: implications for mate choice and sperm competition. Herpetologica 54, 388–394. 10.2307/3893157

56. Tourmente, M., Giojalas, L. C. and Chiaraviglio, M. (2011). Sperm parameters associated with reproductive ecology in two snake species. Herpetologica 67, 58–70. 10.1655/HERPETOLOGICA-D-10-00052.1

57. Quintero-Pérez, R. I., Méndez-de la Cruz, F. R., Miles, D. B., Vera Chávez, M. C., López-Ramírez, Y., Arenas-Moreno, D. M., and Arenas-Ríos, E. (2023). Trade-off between thermal preference and sperm maturation in a montane lizard. J. Therm. Biol. 113, 103526. 10.1016/j.jtherbio.2023.103526

58. van Heerwaarden, B. and Sgrò, C. M. (2021). Male fertility thermal limits predict vulnerability to climate warming. Nat. Commun. 12, 2214. 10.1038/s41467-021-22546-w

59. Vasudeva, R., Dickinson, M., Sutter, A., Powell, S., Sales, K. and Gage, M. J. G. (2021). Facultative polyandry protects females from compromised male fertility caused by heatwave conditions. Anim. Behav. 178, 37–48. 10.1016/j.anbehav.2021.05.016

60. Vasudeva, R., Sutter, A., Sales, K., Dickinson, M. E., Lumley, A. J. and Gage, M. J. G. (2019). Adaptive thermal plasticity enhances sperm and egg performance in a model insect. Elife 8, 1–21. 10.7554/eLife.49452

61. Vervust, B., Lailvaux, S.P., Grbac, I. and Van Damme, R. (2008). Do morphological condition indices predict locomotor performance in the lizard *Podarcis sicula*? Acta Oecol. 34, 244–251. 10.1016/j.actao.2008.05.012.

62. Walsh, B. S., Parratt, S. R., Hoffmann, A. A., Atkinson, D., Snook, R. R., Bretman, A. and Price, T. A. R. (2019). The impact of climate change on fertility. Trends Ecol. Evol. 34, 249–259. 10.1016/j.tree.2018.12.002

63. Walsh, B. S., Parratt, S. R., Snook, R. R., Bretman, A., Atkinson, D. and Price, T. A. R. (2022). Female fruit flies cannot protect stored sperm from high temperature damage. J. Therma. Biol. 105, 103209. 10.1016/j.jtherbio.2022.103209

64. Wang, W. W.-Y., Gunderson A. R. (2022). The physiological and evolutionary ecology of sperm thermal performance. Front. Physiol. 13, 754830. 10.3389/fphys.2022.754830

65. Weimer, M., Jiang, X., Ponta, O., Stanzel, S., Freyberger, A. and Kopp-Schneider, A. (2012). The impact of data transformations on concentration-response modeling. Toxicol. Lett. 213, 292–298. 10.1016/j.toxlet.2012.07.012

66. Williams, S. E., Shoo, L. P., Isaac, J. L., Hoffmann, A. A. and Langham, G. (2008). Towards an integrated framework for assessing the vulnerability of species to climate change. PLoS Biol. 6, 2621–2626. doi:10.1371/journal.pbio.0060325

67. Wolak, M. E., Fairbairn, D. J. and Paulsen, Y. R. (2012). Guidelines for estimating repeatability. J. Mol. Biol. 3, 129–137. 10.1111/j.2041-210X.2011.00125.x

